## Supplementary figures and images for "The relationship of mRNA with protein expression in CD8^+^ T cells associates with gene class and gene characteristics"

### Supplemental Figure 1

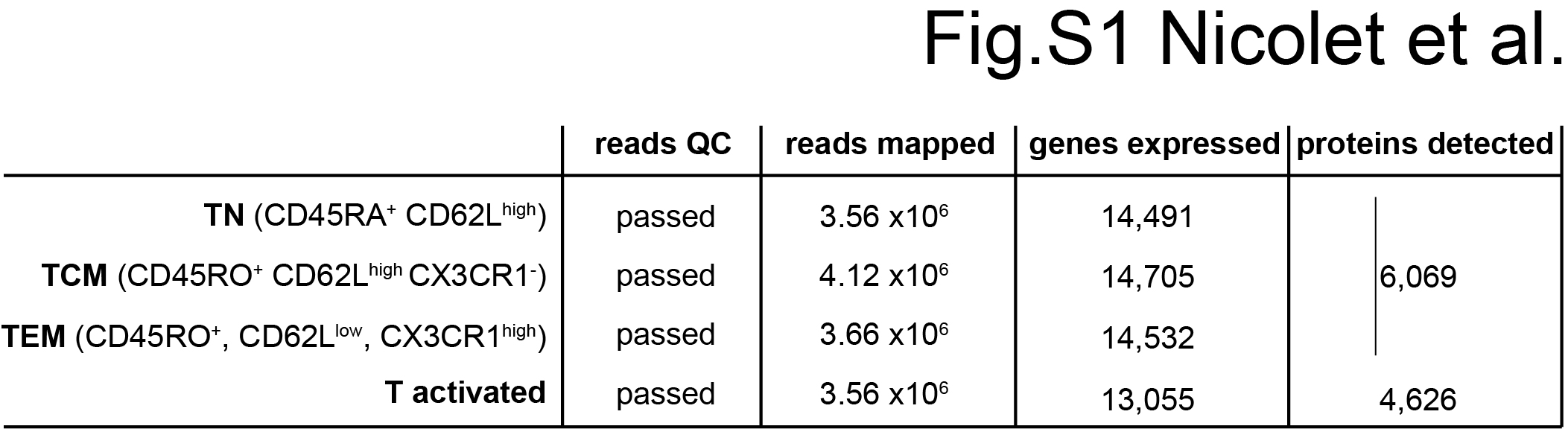

### Supplemental Figure 2

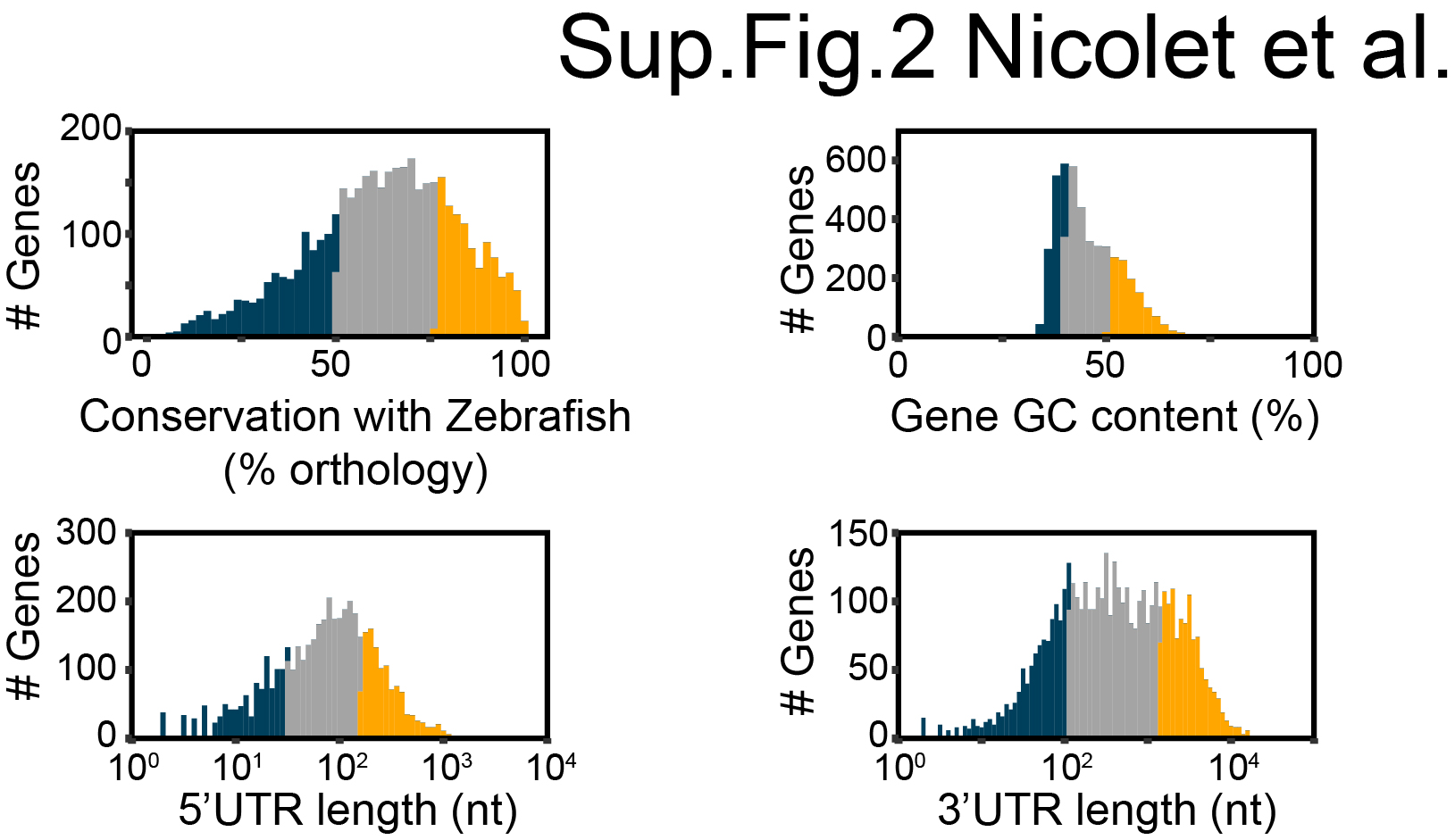

### Supplemental Figure 3

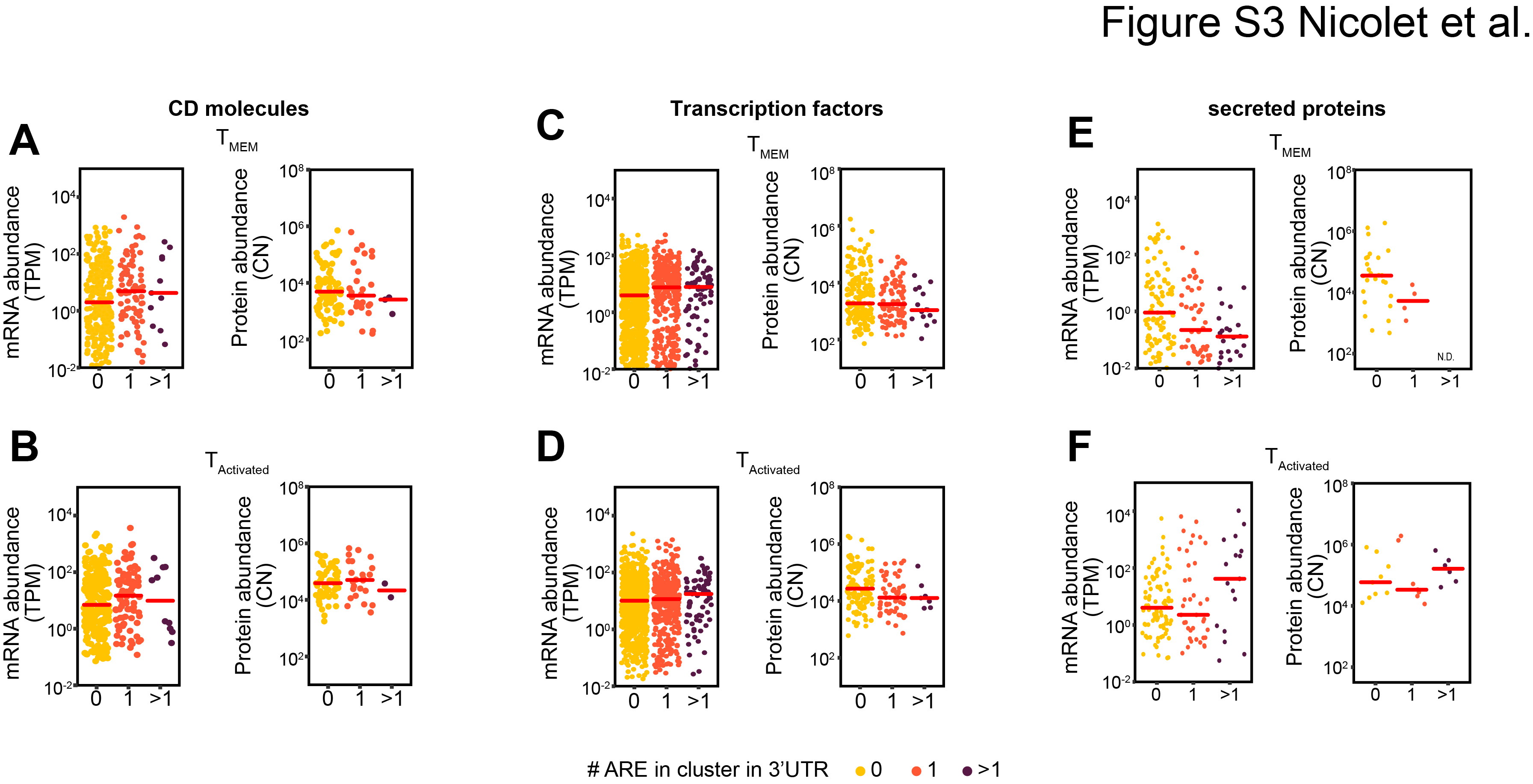
